# Extreme tolerance of *Paramecium* to acute injury induced by γ rays

**DOI:** 10.1101/681098

**Authors:** Ye Wang, Xiaopeng Duan, Yiping Hu

## Abstract

Due to the wide use of nuclear energy, nuclear radiation threat is increasing day by day. However, methods for protection against nuclear radiation are limited and not effective. Therefore, it is necessary to develop an effective method for protection and therapy against injury caused by radiation. According to the law of biological evolution, one of the most primitive eukaryotes, *Paramecium*, could be tolerant to ionizing radiations, in a way similar to or better than *Ramazzottius varieornatus* and *Bdelloids rotifer*. Therefore, in the present study, the anti-radiation effects of *Paramecium* were explored, which could be applicable to human beings. *Paramecium* was found to be tolerant to ^60^Co γ rays, and its half lethal dose (LD50) was approximately 3000 Gy, which is 60 times more than that of human beings. The GSPATT00021409001 and GSPATT00016349001 genes were significantly up-regulated, which might play a role in DNA binding and nuclear transport, and thus, could function in DNA protection or repair. The biological function of the two genes and their effects in humans need to be explored in future studies.

## Introduction

With the development of physics, nuclear energy was discovered, and it presently has multiple applications. As fossil fuel reserves are limited, nuclear energy is expected to become the major source of energy^1^. In recent years, the number of nuclear power stations have increased. Moreover, the storage of nuclear weapons is increasing, and consequently, there is a rise in nuclear weapon threat. Thus, the threat of nuclear radiation damage to human beings has also increased. Although the effects and mechanism of nuclear radiation have been extensively studied^2^, effective prevention and control measures against exposure to nuclear radiation are still unknown.

When life first appeared on the earth, the environmental condition of the prima earth was extreme, as it included high intensity cosmic rays, high temperature, and frequent lightning. Early biological species survived and propagated under extreme conditions, which indicates that the early biological species possessed the mechanisms to tolerate such unfavorable conditions. The radiation intensity on the prima earth then, would have been much higher than it is at present. Thus, some primitive organisms may be more tolerant to nuclear radiation. Researchers have found that *Ramazzottius varieornatus*^3,4^ and *Bdelloids rotifer*^5,6^ are highly tolerant to nuclear radiation, compared to the human beings. Similarly, there may be a few more primitive species, which may also have a higher tolerance to radiation. *Paramecium*, a unicellular organism, is more primitive than *Ramazzottius varieornatus* and *Bdelloids rotifer*. Therefore, the radiation tolerance ability of *Paramecium* needs to be studied.

This study focuses on understanding the effects of ^60^Co γ rays on *Paramecium* and the associated changes in gene expression. The *Paramecium* was irradiated with serial doses of γ ray ranging from 100 Gy to 4000 Gy. Interestingly, *Paramecium* was found to be tolerant to γ ray irradiation, and the half lethal dose (LD50) was approximately 3000 Gy, which is 60 times more than that of human beings. Expression profile sequencing was performed to study the radiation protection genes of *Paramecium*, and significant changes were found in the genes. Thus, these genes may play a role in protection against radiation. Moreover, these genes can also be used to develop new drugs or methods for protection or therapy against radiation injury.

## Methods

### Culturing and isolation of *Paramecium*

A sample from the field was collected, and a single *Paramecium* cell was isolated using a 10 μl micropipette by viewing it under an inverted phase contrast microscope. The single cell was then inoculated into wheat grain medium and incubated at 28°C with normal air.

The wheat grain medium was prepared by using high pressure to treat the wheat grains, followed by drying them in a drying box to improve its shelf life. The treated wheat grains were then boiled in the water for at least 30 min. It was allowed to cool and then exposed to air for 24 h, following which it was used for extended culture of the isolated *Paramecium*.

### Treatment of *Paramecium* using γ rays

*Paramecium* was irradiated with 100 Gy, 500 Gy, 1000 Gy, 2000 Gy, 3000 Gy, and 4000 Gy doses of γ rays. After irradiation, the *Paramecium* was observed and counted under the light microscope at 40× magnification for 5 sec under each visual field. This was repeated 4 times for each dose of radiation. The γ rays were produced from the nuclear decay of ^60^Co, which was provided by the Radiation Center of Second Military Medical University.

### Expression profile sequencing of *Paramecium* treated with γ rays

To explore the mechanism of γ ray tolerance in *Paramecium*, expression profile sequencing was performed. For expression profile sequencing, the *Paramecium* was irradiated with ^60^Co γ rays at a dose of 2000 Gy. After irradiation, the cells were centrifuged and dissolved in RNAiso for further RNA extraction. Total RNA (1 μg) was used for extracting rRNA using the Ribo-Zero rRNA Removal Kit (Illumina, San Diego, CA, USA) according to the manufacturer’s instructions. RNA libraries were constructed by using the RNA devoid of rRNA with the help of the TruSeq Stranded Total RNA Library Prep Kit (Illumina, San Diego, CA, USA), according to the manufacturer’s instructions. Library quality control and quantification was performed using the BioAnalyzer 2100 system (Agilent Technologies, Inc., USA). The library (10 pM) was denatured into single-stranded DNA molecules, captured on Illumina flow cells, amplified *in situ* as clusters and sequenced for 150 cycles on the Illumina HiSeq Sequencer according to the manufacturer’s instructions. High throughput sequencing service was provided by CloudSeq Biotech (Shanghai, China).

### Prediction of the structure and function of different expression genes

Expression profile sequencing of the significant expression genes was carried out, and the structure of the translated proteins was predicted using the SWISS-MODEL (https://www.swissmodel.expasy.org/). The SWISS-MODEL is an automated system for modelling the 3D structure of a protein from its amino acid sequence using homology modelling techniques^7^. The translated proteins was also classified using the ProtoNet 6.0 (http://www.protonet.cs.huji.ac.il) tool, which is a data structure of protein families that cover the protein sequence space.^8^

### Statistical analysis

All data are presented as mean ± SD. Statistical computations were performed using the GraphPad Prism version 6.0. A P<0.05 was considered statistically significant. (*** indicate P<0.001)

## Results and Discussion

### *Paramecium* was highly tolerant to ^60^Co γ rays

*Paramecium* was irradiated with ^60^Co γ rays ranging from 100 Gy to 4000 Gy. After irradiation, the survival situation was observed and the number of *Paramecium* was counted. It was found that the amount of *Paramecium* did not decrease under an irradiation of 2000 Gy. However, the number of *Paramecium* reduced significantly above an irradiation of 3000 Gy. At 4000 Gy, most of the *Paramecium* cells died; however, the few that survived had structural deformity. According to these results, the LD50 (50% lethal dose) of γ rays to *Paramecium* was determined to be approximately 3000 Gy. (Figure 1–2, Supplementary Movie 1-7).

**Figure 1.**
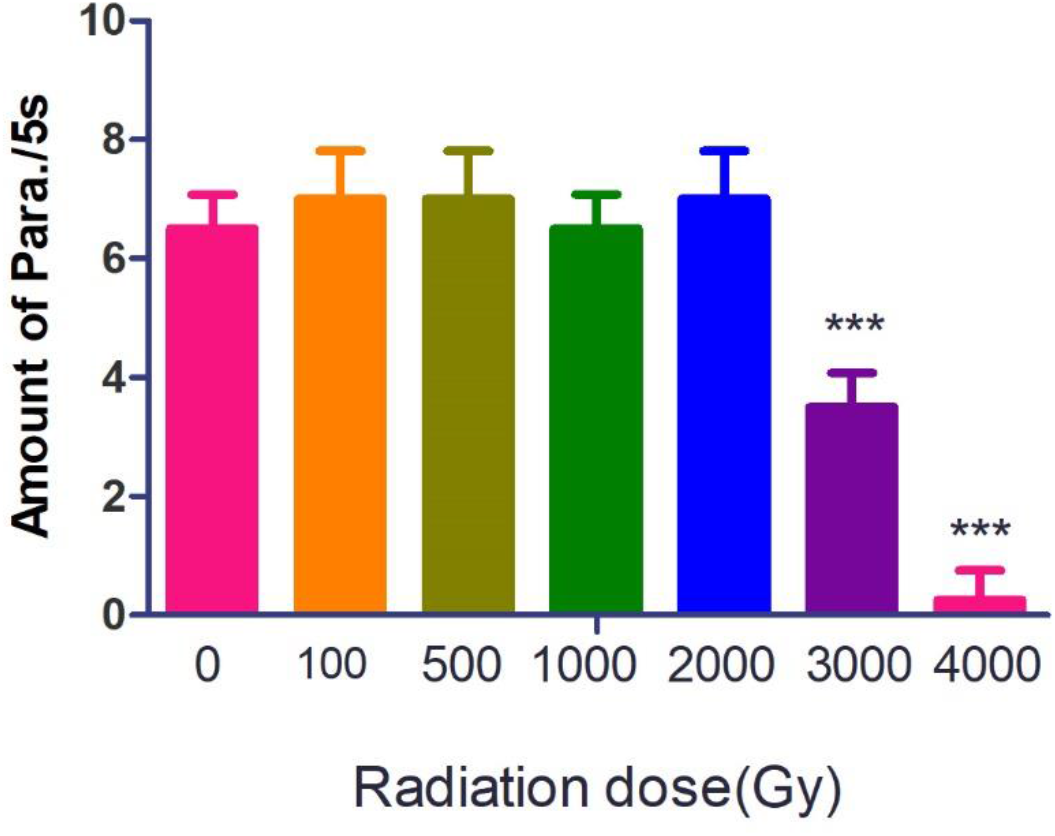
Sensitivity of *Paramecium* to γ rays.

**Figure 2.**
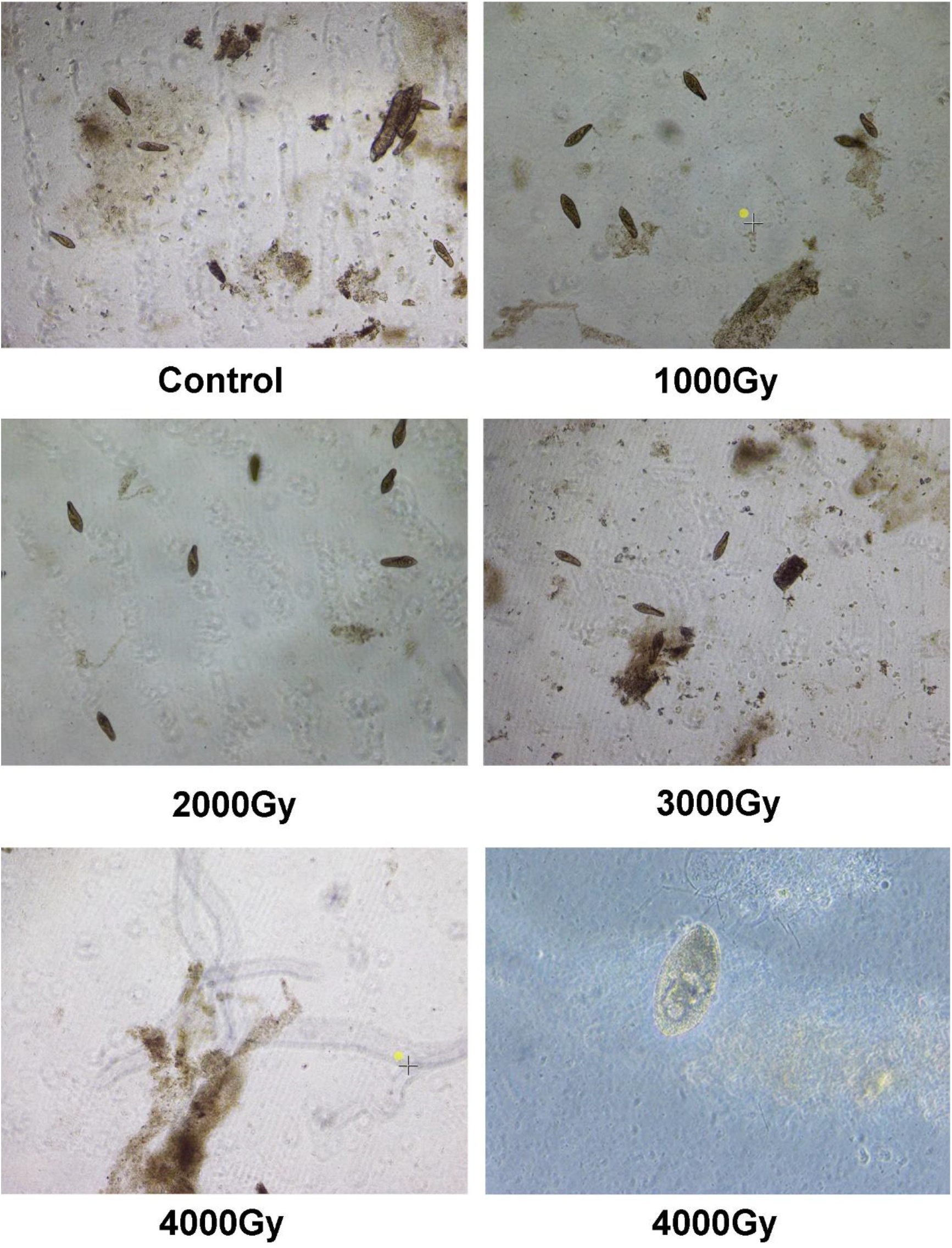
Images of *Paramecium* exposed to different dose of γ rays. (40x magnification and the last image was observed at a magnification of 100x)

A death rate of 100% was observed in human beings exposed to an irradiation dose of over 8 Gy ^9^. An irradiation dose of more than 15Gy, causes acute radiation sickness, leading to immediate nausea and vomiting, and death within 48 h ^9,10^.

However, primitive organisms, such as *Ramazzottius varieornatus* and *Bdelloids rotifer*, are more tolerant to the irradiation than human beings, including whole organism body and cells. Both *Ramazzottius varieornatus* and *Bdelloids rotifer* are highly tolerant to ionizing radiations over a dose of kGy, and the anhydrobiotic condition was more radioresistant than the hydrated one. Thus, according to our results, *Paramecium* showed higher radio-resistance than human beings, similar to *Ramazzottius varieornatus* and *Bdelloids rotifer*. There might be a common mechanism used by these primitive species to tolerate the radiations which needs to be explored.

### γ rays induced high expression genes may play a role in tolerance towards radiation

Studying the gene expression changes of *Paramecium* irradiated with γ rays will help understand the radiation protection mechanism of *Paramecium*. By using high-throughput sequencing methods, the expression profile sequencing of *Paramecium* treated with γ rays at a dose of 2000 Gy was studied. It was found that exposure to γ rays up-regulated expression of 25 genes and down-regulated expression of 34 genes in *Paramecium*. A p value lower than 0.05 was obtained (Figure 3A, B).

**Figure 3.**
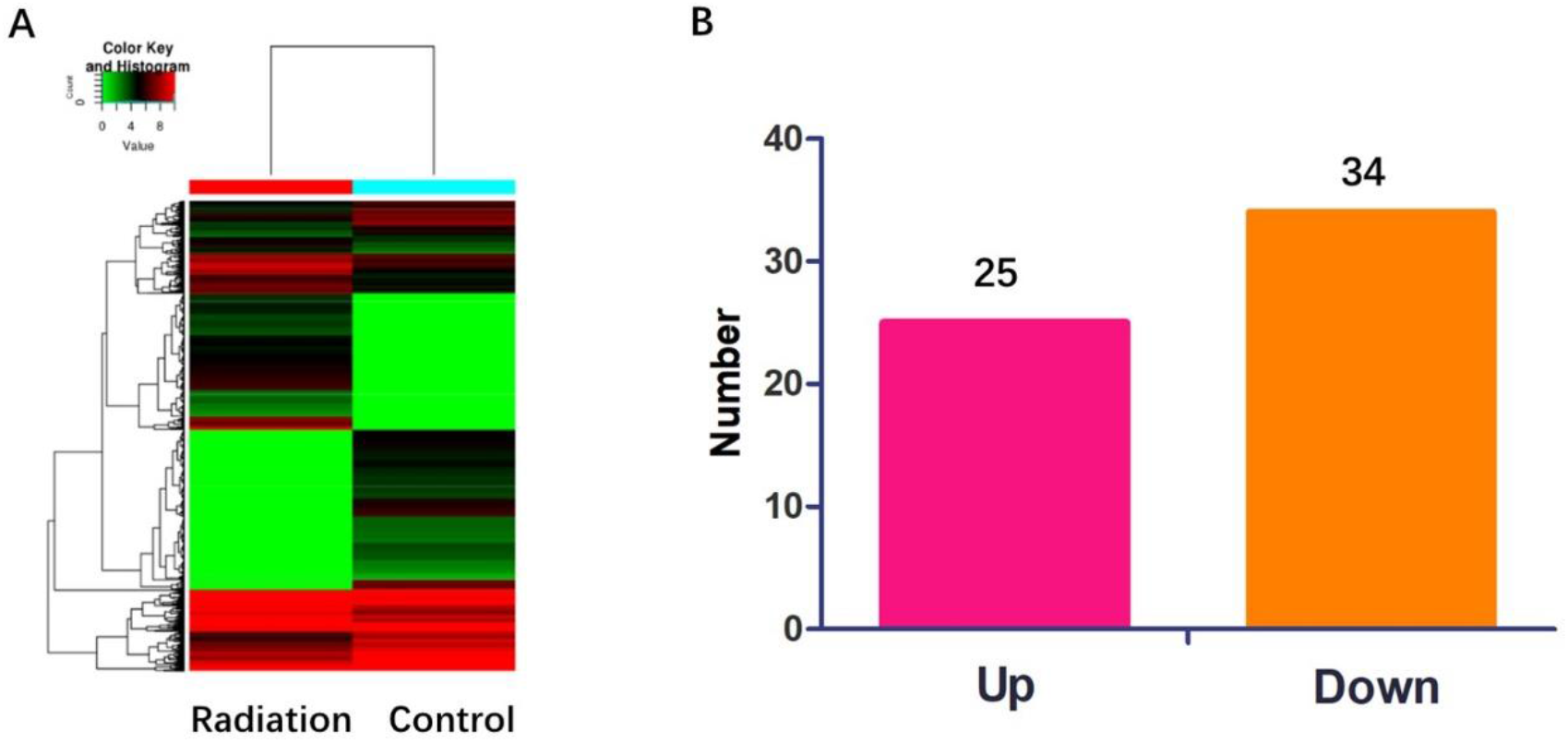
(A) The heatmap of the expression profile sequencing (B) The number of changed expression genes.

Among the upregulated genes, the three genes that were strongly up-regulated, were gene IDs GSPATT00021409001, GSPATT00016349001, and GSPATT00004424001 (Supplementary Table 1). These three genes were protein coding genes. Since proteins play an importance role, it was necessary to predict the function of the three upregulated genes. Using the SWISS-Model and ProtoNet, it was found that GSPATT00021409001 gene expressed proteins that may form a homo-tetramer, and showed a DNA/RNA binding function (Figure 4A), which indicates that GSPATT00021409001 protein may target the DNA after being irradiated and may have DNA protective effects, similar to the Dsup protein of Tardigrades. Moreover, GSPATT00016349001 protein was predicted to form a homo-dimer, and it may be a member of the nuclear transport factor 2 (Figure 4B), which may have the function of carrying the protein through the nucleopore, surely including the GSPATT00021409001 protein. The GSPATT00021409001 and GSPATT00016349001 proteins were both produced in response to the radiation, and were up-regulated significantly. GSPATT00021409001 protein was carried by GSPATT00016349001 protein into the nucleus, and enabled binding to the DNA of *Paramecium*, wherein it performed the DNA protection or repair function for the survival of *Paramecium*. GSPATT00021409001 and GSPATT00016349001 proteins may function together to confer radiation resistance to the *Paramecium*. The biological function of these two proteins will be explored further. Furthermore, the structure and function of GSPATT00004424001 could not be predicted by the SWISS-Model, which also needs to be explored in future studies.

**Figure 4.**
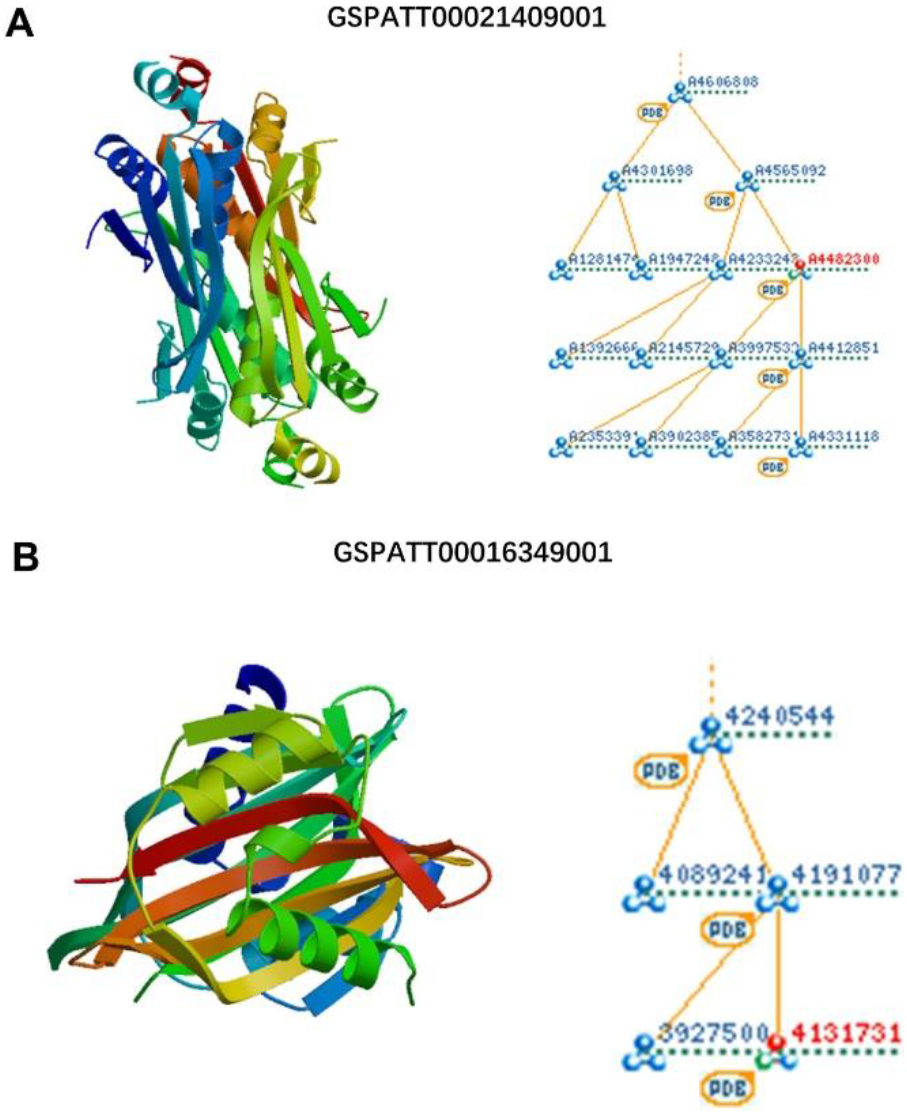
The predictive protein 3D structure and classification of GSPATT00021409001 and GSPATT00016349001 genes.

## Conclusion

In this study, we showed that *Paramecium* was strongly tolerant to ^60^Co γ rays, and the LD50 was approximately 3000 Gy, which was much higher than the LD50 of human beings. Expression profile sequencing assay showed that γ rays can induce various gene expression changes. Among these, two upregulated genes, including GSPATT00021409001 and GSPATT00016349001 genes, may perform important functions in anti-radiation damage, according to the function prediction tool. However, certain biological effects of GSPATT00021409001 and GSPATT00016349001 proteins are still unknown. Moreover, the ability of the γ ray protection genes of *Paramecium* to replicate their effects in human cells needs to be further studied.

